# A causal analysis of the effect of age and sex differences on brain atrophy in the elderly brain

**DOI:** 10.1101/2020.11.20.391623

**Authors:** Jaime Gómez-Ramírez, Miguel A. Fernández-Blázquez, Javier González-Rosa

## Abstract

We study how brain volume loss at old age is affected by factors such as age, APOE gene, sex, and school level. The quantitative characterization of brain volume loss at old age relative to young age requires at least in principle two MRI scans performed at both young and old age. There is, however, a way to address the problem by having only one MRI scan at old age. We compute the total brain loss of elderly subjects as the ratio between the estimated brain volume and the estimated total intracranial volume. Magnetic resonance imaging (MRI) scans of 890 healthy subjects aged 70 to 85 were assessed. The causal analysis of factors affecting brain atrophy was performed using Probabilistic Bayesian Modeling and the Mathematics of Causal Inference. We find that healthy subjects get into their seventies with an average brain volume loss of 30% from their maximum brain volume at a young age. Both age and the sexes are causally related to brain atrophy, with women getting to elderly age with 1% larger brain volume relative to intracranial volume than men. How the brain ages and what are the reasons for sex differences in adult lifespan are causal questions that need to be addressed with causal inference and empirical data. The graphical causal modeling presented here can be instrumental in understanding a puzzling scientific inquiry -the biological age of the brain.

## I. Introduction

The historical investigation of brain volume variation with age has at least three well defined periods: the era of autopsies, followed by the utilization of magnetic resonance imaging (MRI) to the present time dominated by computational anatomy making use of MRI leveraged by data-analytical methods.

Early evidence of the effect of aging on brain size and structure comes from autopsy studies in the XIX century that indicated that brain weight reduced surely but slowly with age [Boyd, 1861], [Marshall, 1892]. Autopsies helped to solidify the commonly held belief that brain weight is stable between the ages of 20 and 50 to progressively decay thereafter. Large sample autopsy-based studies, still prior to the MRI era, suggest that brain weight reaches its maximum in the late teens and declines very slowly (0.1 0.2% a year) till the age of 60s-70s, after that the decline is faster [Miller and Corsellis, 1977],[Esiri, 2007]. In a 1980 study [Ho et al., 1980], weights of fresh brains of 1,261 subjects aged 25 to 80, showed that the brain mass decreases rapidly after age 80. It also indicated different atrophy patterns based on ethnicity and sex: *“the rate of decrease for the brain weight after age 25 years is highest for white men, followed by black women, white women, and black men, and, except that between white men and white women, the differences are statistically insignificant.”* Around the same time, brain autopsies indicated that progressive decline in brain weight begins at about 45 to 50 years of age to reach its lowest values after the age of 86 [Dekaban and Sadowsky, 1978],. The study postulated that the maximum brain weight attained in young adults was reached at 19 years of age, estimating an accumulated loss of brain weight of 11% between the ages 19 and 86 and detecting differential rates of change in brain weight depending on age and less so by sex. However, studies based on autopsies present problems of reliability, selection bias and most importantly, they can’t tell us anything about cerebral atrophy in living individuals. The advent of non-invasive imaging changed this.

Magnetic Resonance Imaging and before that computed tomography, created the possibility of assessing cerebral volume in-vivo, non-invasively, and repeatedly [Fox and Schott, 2004]. Imaging studies revealed global volume loss and regional variation as major effects of aging in the brain. Nevertheless, the estimates of volume and tissue loss were prone to error because they required manual outlining and furthermore included a strong bias since the brain areas are selected a priori [Wenger et al., 2014], [Despotović et al., 2015].

The advent of new computerized methods sensitive to variations in size, shape, and tissue characteristics of brain structures, set the final stage in the study of brain anatomy in aging, offering a new set of tools unknown to previous researchers who needed to rely upon autopsies and manual outlining of MRI and tomographies [Ashburner et al., 2003]. Specifically, the game-changer event was Voxel-Based-Morphometry (VBM), a whole-brain, an unbiased technique for characterizing regional cerebral volume and tissue concentration differences in structural magnetic resonance images. MRI studies with automatic segmentation of in vivo aging brains have proliferated since then. In [Good et al., 2001], a tissue-based segmentation study of the effects of aging on gray and white matter and cerebral spinal fluid (CSF) in 465 normal adults, showed that gray matter volume decreased linearly with age, with a significantly steeper decline in males. Global white matter, on the other hand, did not decline with age. In addition to this, the study found that brain aging behaves locally, from areas with accelerated loss such as the insula, superior parietal gyri, central sulci, and cingulate sulci to areas with little age effect, notably the amygdala, hippocampi, and entorhinal cortex. Sowell and colleagues [Sowell et al., 2003], in a study of 176 normal individuals ranging in age from 7 to 87 years, found that areas known to myelinate early showed a more linear pattern of aging atrophy than the frontal and parietal neocortices, which continue myelination into adulthood. Along these lines, the study suggested that areas involved in language functions such as the left posterior temporal cortices, have a more protracted course of maturation than any other cortical region.

There is growing evidence that age imposes a stronger influence on brain structure in older than in younger adults, but the onset and the type of decline (linear, non-linear) depends on tissue and brain region. The common understanding of tissue atrophy points to gray matter atrophy onset may start in young adulthood, around 18, white matter, on the other hand, remains relatively stable until old age. Although we are lacking a theory of human brain aging capable of making robust predictions about brain growth and atrophy, we know that rapid growth occurs during childhood/adolescence, with a particularly dramatic growth rate during the first 3 months, approximately 1%per day, reaching the half of elderly adult brain volume by the end of the first 3 months [Holland et al., 2014]. Between 18 and 35 years old, the brain experience a period of consolidation with no brain tissue loss. After 35 years, Hedman and colleagues [Hedman et al., 2012] have suggested a steady volume loss of 0.2% per year, which accelerates gradually to an annual brain volume loss of 0.5% at age 60. After 60 years of age, the same study indicates a steady volume loss of more than 0.5%

Fjell et al. [Fjell and Walhovd, 2010] found nonlinear decline across chronological age in the hippocampus, caudate but linear decline slopes for thalamus and accumbens. In [Rast et al., 2017], cortical thinning was found to be significantly altered by hypertension and Apolipoprotein-Ee4 (APOEe4), with frontal and cingulate cortices thinning more rapidly in APOEe4 carriers. Additional longitudinal studies have found different brain atrophy patterns according to clinical conditions, including cognitive decline and Alzheimer’s disease [Rusinek et al., 2003], [Chételat et al., 2005], [Misra et al., 2009] and multiple sclerosis [Ghione et al., 2019] to cite a few. Nonetheless, small sample size and the lower reliability for segmenting small structures are important caveats in longitudinal studies [Oschwald et al., 2019].

There is, however, an approach that to our knowledge has not been undertaken with the sufficient sample size and the proper methodology. We are referring to quantify the brain loss at older age relative to the brain’s maximum size reached at some point in her young age. Although the global brain volume lacks the required granularity to map onto its cognitive processes of interest such as memory or language, it posses, on the other hand, the singular property that it is possible to infer approximately its maximum volume via the intracranial volume (ICV). The ICV acts as a scaffolding of the brain and sets an upper bound for the brain’s volume. Accordingly, it is possible to build a proxy of the brain atrophy that an elder person went through her adult life by means of computing the ratio between the brain volume (BV) estimation at the moment of the MRI scan and the ICV which represents the upper limit of brain volume. Thus, 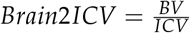.

By quantifying, even if in an approximative way, the brain volume loss at older age relative to its upper bound at a young age (Brain2ICV), we can make educated guesses about the effect of brain aging in a person. The mismatch between the actual brain volume and the expected brain volume according to the person’s age might contain valuable information to better understand brain aging dynamics.

## II. Methods

The dataset used here comes from a single-center, observational cohort study of 1,213 subjects [Gómez-Ramírez et al., 2019], [Fernández-Blázquez et al., 2020]. The participants of the study are home-dwelling elderly volunteers, aged 69 to 85, without relevant psychiatric, neurological, or systemic disorders. Of the initial 1,213 subjects, those that were diagnosed with MCI or dementia plus those lacking a brain MRI were excluded from our analysis, resulting in a cohort of 890 healthy elderly subjects. After signing informed consent, the participants undertake a yearly systematic clinical assessment including medical history, neurological and neuropsychological exam, blood collection and brain MRI. Ethical approval was granted by the Research Ethics Committee of *Instituto de Salud Carlos III* and written informed consent was obtained from all the participants. The authors assert that all procedures contributing to this work comply with the ethical standards of the relevant national and institutional committees on human experimentation, and with the Helsinki Declaration of 1975 and its later amendments.

The level of education was encoded as 0 *no formal education*, 1 *primary education*, 2 *middle or high school degree* and 3 *university degree*. Cognitive status was determined with the Mini-Mental Status Examination (MMSE), Free and Cued Selective Reminding Test (FCSRT), Semantic fluency, Digit-Symbol Test and Functional Activities Questionnaire (FAQ). APOE genotype was studied with total DNA isolated from peripheral blood following standard procedures. The APOE variable was coded 1 for the *e*4-carriers, and 0 for non-carriers. Family history of AD was coded as 0 for subjects with no parents or siblings diagnosed with dementia and 1 for those with at least one parent or sibling diagnosed with dementia.

The imaging data were acquired in the sagittal plane on a 3T General Electric scanner (GE Milwaukee, WI) utilizing T1-weighted inversion recovery, supine position, flip angle 12°, 3-D pulse sequence: echo time *Min. full*, time inversion 600 ms., Receiver Bandwidth 19.23 kHz, field of view = 24.0 cm, slice thickness 1 mm and Freq × Phase × 288 × 288. The brain volume loss at the moment of having the MRI compared to the maximum brain volume is computed as the Brain Segmentation Volume to estimated Total Intracranial Volume (eTIV) [estimated Total Intracranial Volume aka ICV, 2020] ratio (ICV and eTIV the FreeSurfer term for intracranial volume are used equivalently). The postprocessing was performed with FreeSurfer [Fischl, 2012], version freesurfer-darwin-OSX-ElCapitan-dev-20190328-6241d26 running under a Mac OS X, product version 10.14.5. For the sake of illustration, Figure 1 shows the result produced of the intracranial volume segmentation for two subjects in the study.

**Figure 1:**
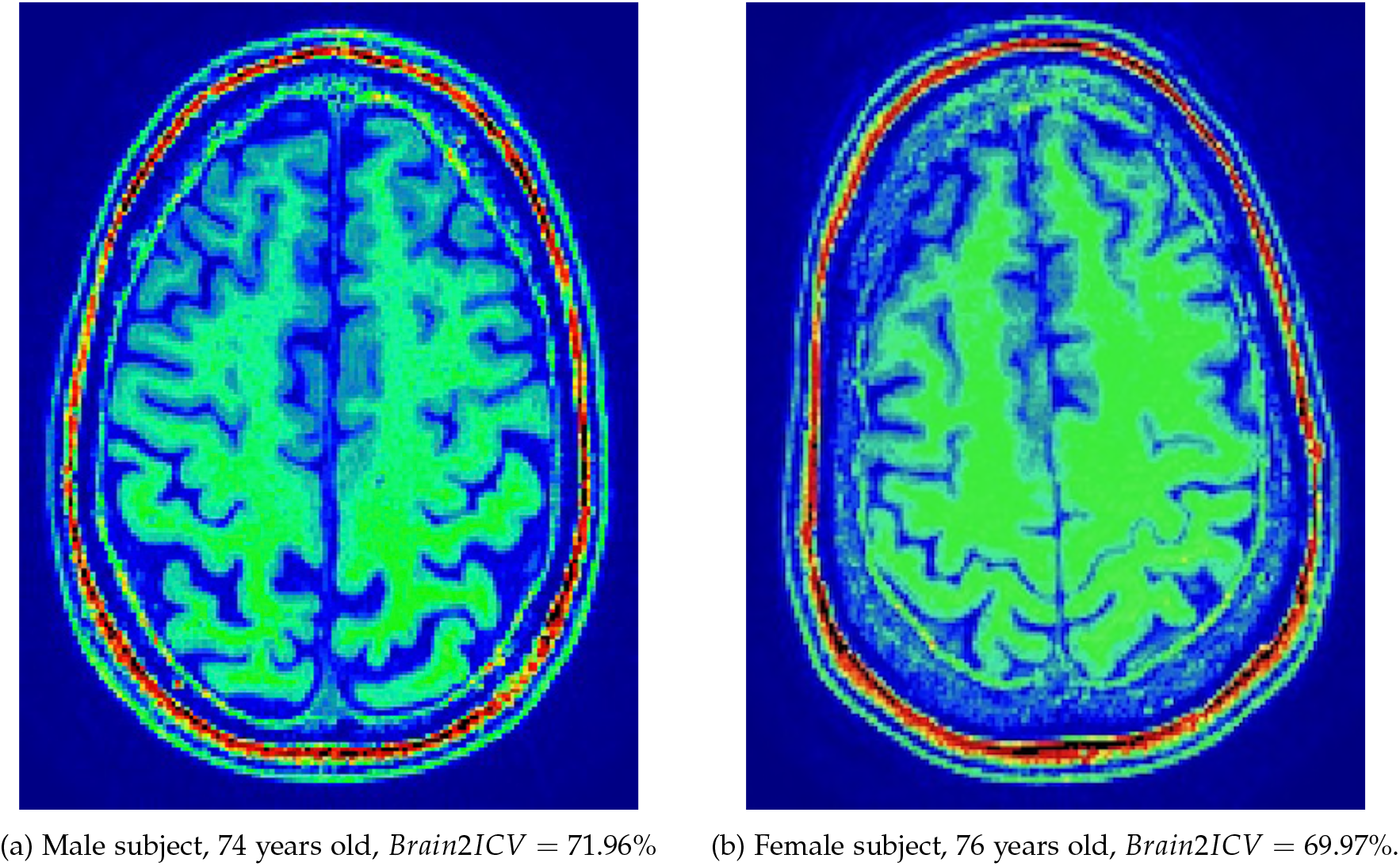
Figure 1a and Figure 1b show an axial view of the brain and its membranous envelopes of two subjects in the study. The Brain2ICV or brain volume to intracranial volume ratio is *Brain*2*ICV* = *TBV*/*eTIV* = 929035/1460465 = 0.7196 for left figure, and *Brain*2*ICV* = *TBV*/*eTIV* = 929035/1327593 = 0.6997 for the right figure.

The eTIV estimated by FreeSurfer has previously been reported to have linear correlations of 0.9 with manually estimated intracranial volume [Shen et al., 2010], [Malone et al., 2015]. Depending on whether CSF is included or not, one can dissociate the total brain volume (TBV) from the intracranial volume (ICV). The normalized TBV (NBV) is widely used as an index for brain atrophy, as the head size remains stable across the life span and serves as a good measure to reduce between-subject differences with regard to maximum brain size. Whole-brain volume, on the other hand, changes throughout the life span of an individual. Measurements of total brain volume (TBV) with FreeSurfer are robust across field strength [Heinen et al., 2016]. It might be noted that the estimated intracranial volume (eTIV) from FreeSurfer is not segmentation-based but calculated from the alignment to the MNI305 brain atlas [Klasson et al., 2018]. FreeSurfer exploits the relationship between the intracranial volume and the linear transform to MNI305 space rather than counting pixels inside the cranium which would be prone to errors because the skull and the Cerebral Spinal Fluid are both dark on a T1 image [Buckner et al., 2004].

Table 1 includes the description of the variables considered in this study, providing the mean and the standard deviation for the continuous variables -Age, Memory Test Score and the brain volume to intracranial volume ratio (Brain2ICV)- and the classes together with the number of elements for each class for the categorical variables -Sex, APOE, Family history of AD, and School level. In order to assess the strength of the linear association between Brain2ICV and the predictor variables, we perform Pearson’s correlation, point biserial correlation, and analysis of variance depending on whether the variable is continuous, dichotomous as in Sex, Family history of AD, and APOE or discrete with more than two values as in School level.

**Table 1:** Summary of the variables used in the study: Age, Sex, APOE, School Level, Family history of AD, Memory Test Score and the estimated ratio between brain and intracranial volumes (Brain2ICV). The mean and the standard deviation is displayed for the continuous variables and the size of each class for categorical variables.

| Variables | Mean | SD |
| --- | --- | --- |
| Age | 74.72 | 3.86 |
| Memory Score | 9.41 | 2.66 |
| Brain2ICV | 0.70 | 0.03 |
|  | N | % |
| Sex |  |  |
| Males | 303 | 34.04 |
| Females | 587 | 65.96 |
| APOE |  |  |
| Non-carriers | 726 | 81.57 |
| Heterozygous $\epsilon 4$ | 157 | 17.64 |
| Homozygous $\epsilon 4$ | 7 | 0.79 |
| School level |  |  |
| No formal education | 170 | 19.10 |
| Primary education | 265 | 25.17 |
| High School | 224 | 25.17 |
| University | 231 | 25.96 |
| Family history of AD |  |  |
| No | 670 | 75.28 |
| Yes | 220 | 24.72 |

### i. Causal analysis

Correlation is the degree to which two variables show a tendency to vary together. Causality, on the other hand, is about the relationship between an observed effect and what caused it. For variable *C* to cause another variable *E*, (*C* → *E*), there must be a flow of information from the cause *C* to the effect *E*. Here, we want to identify the causal paths built on top of correlation paths that link one or more causes with an effect, specifically the variables that causally affect Brain2ICV. Thus, we aim at studying causal connections between correlated variables using Probabilistic Bayesian Modeling [Davidson-Pilon, 2015] and the mathematics of causal inference, *do-calculus*, proposed by Pearl [Pearl and Mackenzie, 2018].

Bayesian data analysis relies upon generative models that can be used to postulate how the available data was generated. Thus, Bayesian data analysis imposes a model-based approach upon the observed data. Models are the mathematical formulation of the observed events and the model parameters are the factors in the model affecting the observed data.

To determine the evidence for hypothesis H1 versus the alternative null hypothesis H0, we need a model of H0 and a model of H1. While frequentist hypothesis testing is based on the squared error one would expect in many identical repetitions of the experiment, Bayesian inference defies this notion for unrealistic. As a matter of fact, on many occasions, it is not only impractical but impossible to replicate experiments which invalidate the frequentist’s notion of using averages for judging an estimator [Samaniego, 2010], [Benjamin et al., 2018]. Bayesian model selection aims at computing the posterior distribution which contains all the information needed about the model parameters. The posterior distribution also allows us to generate predictions based on actual data and the estimated parameters. Once we have the posterior distribution we can use it to make predictions, *ŷ*, based on data *y* and the estimated parameters, *θ*. The posterior predictive distribution is an average of conditional predictions over the posterior distribution of *θ* (Equation 1).

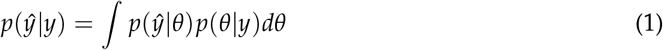

Bayesian Networks are probabilistic graphical models that represent the dependencies of a set of variables and their joint distribution [Pearl, 2009]. Specifically, a Bayesian Network is a graph with directed edges (associations) and with acyclic structure, that is, a node can’t be its own ancestor or descendant. Directed Acyclic Graphs or DAG for short provides a graphical representation of the causal relationships between variables. In a DAG, contrary to statistical models, it is possible to detect the conditional independence between variables.

Figure 2 shows the DAG used to model the causal relationships between the variables in the study. We are particularly interested in clarifying the causal structure in the colored nodes depicted in Figure 2, that is, how Sex and Age affect Brain2ICV. Sex and Age are according to the DAG, the only two relevant variables that directly influence the Brain2ICV. It is worth remarking that a DAG cannot be directly generated from observational data alone, the structure of the DAG makes use of expert knowledge. Once the DAG is in place, it can be used to guide interventions that substantiate the causal reasoning that emanate from the DAG.

**Figure 2:**
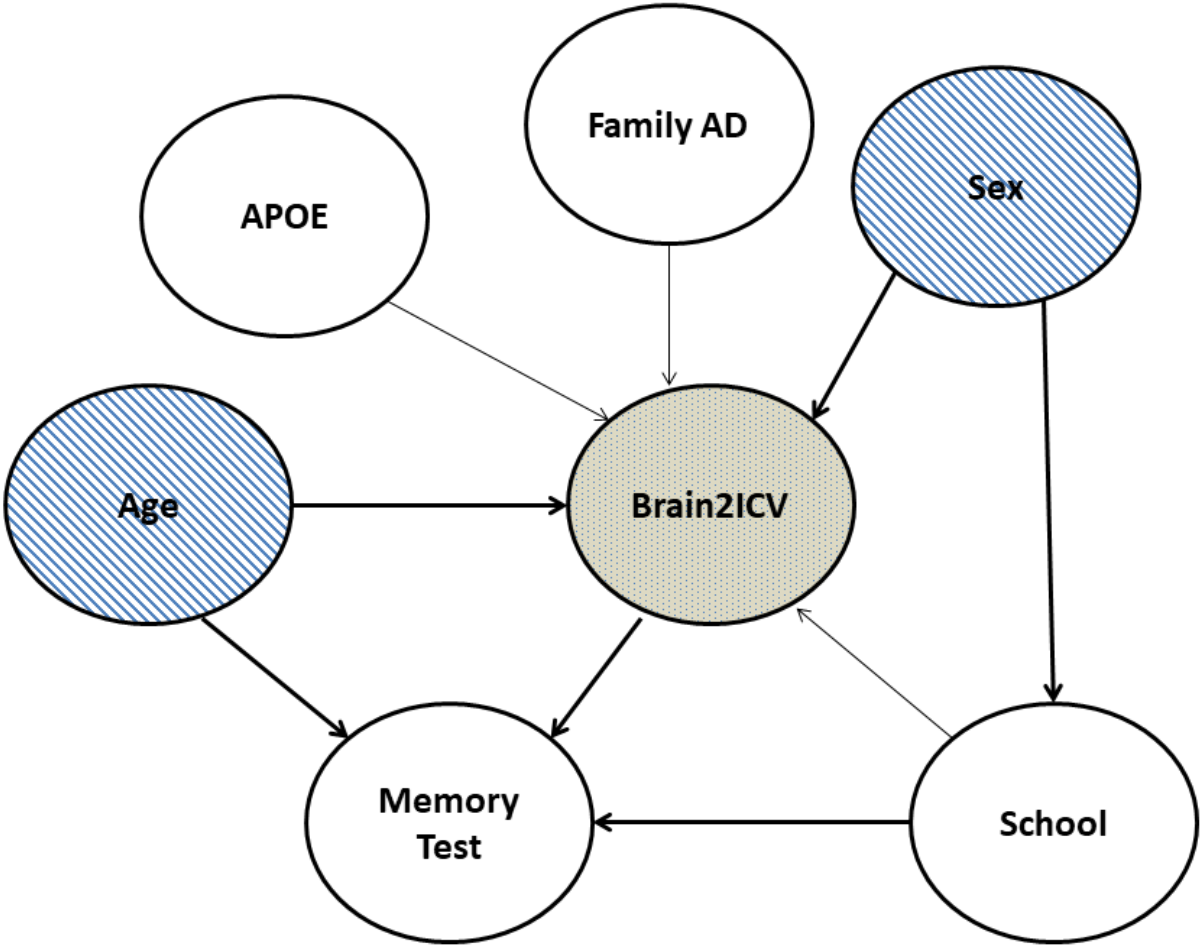
Directed Acyclic Graph (DAG) that postulates possible causal relationships between the variables in the study. A variable C in a causal diagram can only causally affect a variable E when there is a directed path from C to E. Based on the DAG, Sex directly influences Brain2ICV and School, Age directly influences Brain2ICV and Memory and School Level directly influences Memory. The correlation coefficient according to the correlation matrix shown in Figure 4 The strength of the arrows, thin or thick, denotes the correlation coefficient according to the correlation matrix shown in Figure 4.

While the association between age and brain atrophy is indisputable, how sex mediates in brain atrophy is unclear. Furthermore, whether Brain2ICV is associated with one predictor (e.g. Age) after conditioning on the other (e.g. Sex) deserves also attention. As we have previously stated, since the questions are causal they cannot be answered from data alone and a model of the process that generate the data is required, in actuality one model for each question. The questions we need to answer are explicitly stated next.

**Q1: Is there a direct causal relationship between Sex and Brain2ICV?**

**Q2: Is there additional value for Brain2ICV in knowing Sex if we already know Age?**

Let us start with *Q*1, in order to investigate the relationship between Sex and Brain2ICV we build the probabilistic Bayesian model shown in the system of Equations 2.

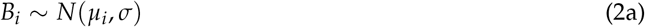

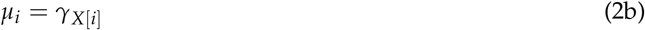

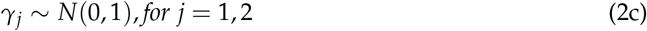

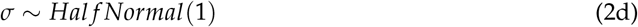

where *B_i_* denotes the variable Brain2ICV or the ratio between brain volume at elderly age and maximum brain volume at a young age for subject *i*, the index variable for Sex *γ*_*j*_, with index *j* = 1, 2, represent the average of Brain2ICV for male (*j* = 1) and female (*j* = 2) (no order implied), which are normally distributed using the same prior, *N*(0, 1), for both male and female subjects. The prior *σ* is assumed to be normally distributed, half-normal to be exact with standard deviation equal to 1 (half-normal distribution can be directly sampled from a Normal distribution by taking the absolute value of each sampled value). Other prior distributions e.g. Exponential or Uniform can also be used.

The last question stated above, Q2, requires a multiple regression model with two predictors Sex and Age. We aim at understanding the following point: Once we know the Sex of an individual, Is there additional predictive power for Brain2ICV in also knowing her Age? And similarly for knowing Sex once we know Age. Thus, we need to quantify each effect and how the three variables are associated finding the conditional independence among them. Formally, *Y* ⫫ *X|Z*, where *Y* is Brain2ICV and *X* is Age (or Sex) and *Z* is Sex or Age. The model that predicts Brain2ICV using both Sex and Age is described in the system of Equations 3.

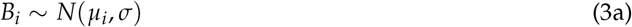

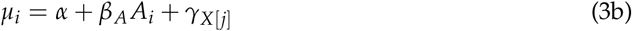

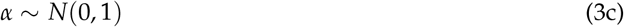

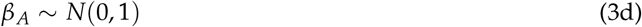

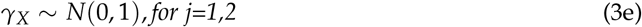

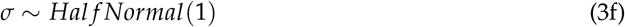

As in the previous equation, *B_i_* denotes the variable Brain2ICV, *γ*_*j*_ represents the average of Brain2ICV for (*j* = 1) male and (*j* = 2) female and the prior distribution of *σ* is half-normal. *A_i_* is the Age of subject *i*. Since all three variables are standardized, we expect the intercept *α* and the parameter *β*_*A*_ to be around zero.

## III. Results

We show first the results of the statistical and correlation analysis in Section i, next in Section ii we describe the causal inference results using probabilistic programming and causal diagrams.

### i. Statistical and Correlation Analysis

Figure 3 shows the violin plot of Age grouped by Sex (Figure 3a) and Brain2ICV grouped by Sex (Figure 3b). Hypothesis testing of the age of males and females gives no difference between the two groups *p* = 0.572 (Figure 3a). On the other hand, the t-test for the means of the Brain2ICV of males and females (Figure 3b) gives a *p* = 2.583^−^8 less than the threshold of 1%, disproving the null hypothesis. According to Figure 3b, females, on average, get to old age (70 or older) with 1.076% less brain atrophy than males as explained by the brain to intracranial volume ratio variable (Brain2ICV). This finding is in agreement with previous work that identified sex differences in the brain during aging and in neurodegenerative diseases. In particular, the thesis that females might have more youthful brains as compared to males is supported by forensic and postmortem studies [Dekaban and Sadowsky, 1978], [Ho et al., 1980]. This hypothesis has been tested very recently in vivo with PET imaging, showing a more persistent metabolic youth in the aging female brain compared to the male brain [Goyal et al., 2019].

**Figure 3:**
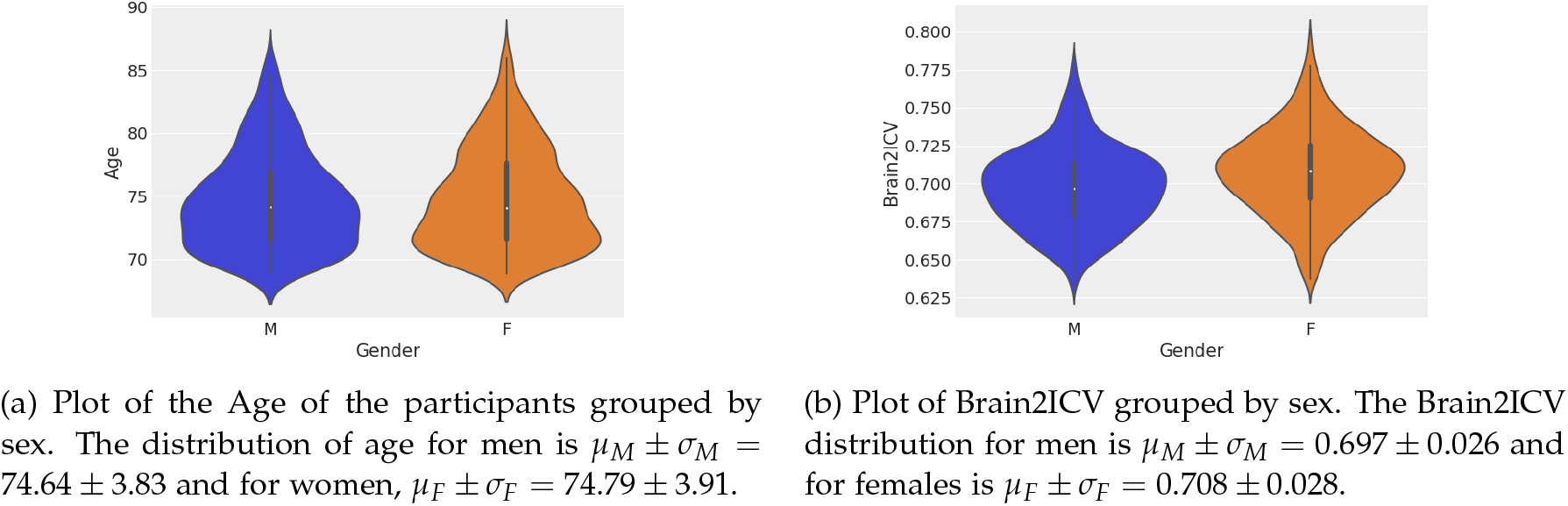
Violin plots of sex distribution (left) and the Brain2ICV grouped by sex (right). The t-test for the means of the two independent samples of scores composed of the age of males and females gives no difference between the groups *p* = 0.572. (Figure 3a). The t-test for the means of the Brain2ICV of males and females gives a p-value 2.583^−8^ *<* 0.01 (Figure 3b).

Table 2 shows the Analysis of Variance (ANOVA) with a linear ordinary least squares (OLS) model [Seabold and Perktold, 2010] which includes the sum of squares, the F statistic and the value of Prob(F) or the probability that the null hypothesis for the null model is true. We are interested in the variables that may have an effect on Brain2ICV which according to the DAG depicted in Figure 2 are Age, APOE, FamilyAD, Sex and School Level. As Table 2 shows, both Age and Sex have a p-value for F statistics less than the significance level of 1%, therefore the null hypothesis -Age, Sex- have no effect on the brain to ICV volume- is therefore rejected. The variable Memory Test is not included since we are interested in variables that potentially cause Brain2ICV, that is to say, the directionality of the arrow must be directed towards Brain2ICV. (The complete summary table of the OLS model *Brain2ICV = Age + Sex + Apoe + School Level + Familial AD* is shown in Supplementary Material, Table 5.)

**Table 2:** Analysis of Variance with a linear OLS Model. Both Age and Sex score show a p-value for F statistic less than the significance level 0.01 to reject the null hypothesis (i.e. age/sex have no effect on the brain to ICV volume). The APOE gene, familial AD and the school level, on the other hand, do not have statistical effect on brain atrophy.

|  | sum sq | F | PR(>F) |
| --- | --- | --- | --- |
| Age | 0.077388 | 119.694321 | **3.242352e-26 |
| Sex | 0.021172 | 32.745601 | **1.437815e-08 |
| APOE | 0.000710 | 1.098909 | 2.947922e-01 |
| Family history of AD | 0.000660 | 1.021550 | 3.124283e-01 |
| School Level | 0.002283 | 3.530481 | 6.057913e-02 |

Before going into the causal analysis results, we study the statistical dependence between every pair of variables in the study. The variables with larger Pearson’s correlation coefficient with Brain2ICV are Age (*ρ* = −.33), Sex (*ρ* = .19) and Memory score test (*ρ* = .14) which are negatively correlated (*ρ* = −.18) among them as the correlation matrix depicted in Figure 4 indicates.

**Figure 4:**
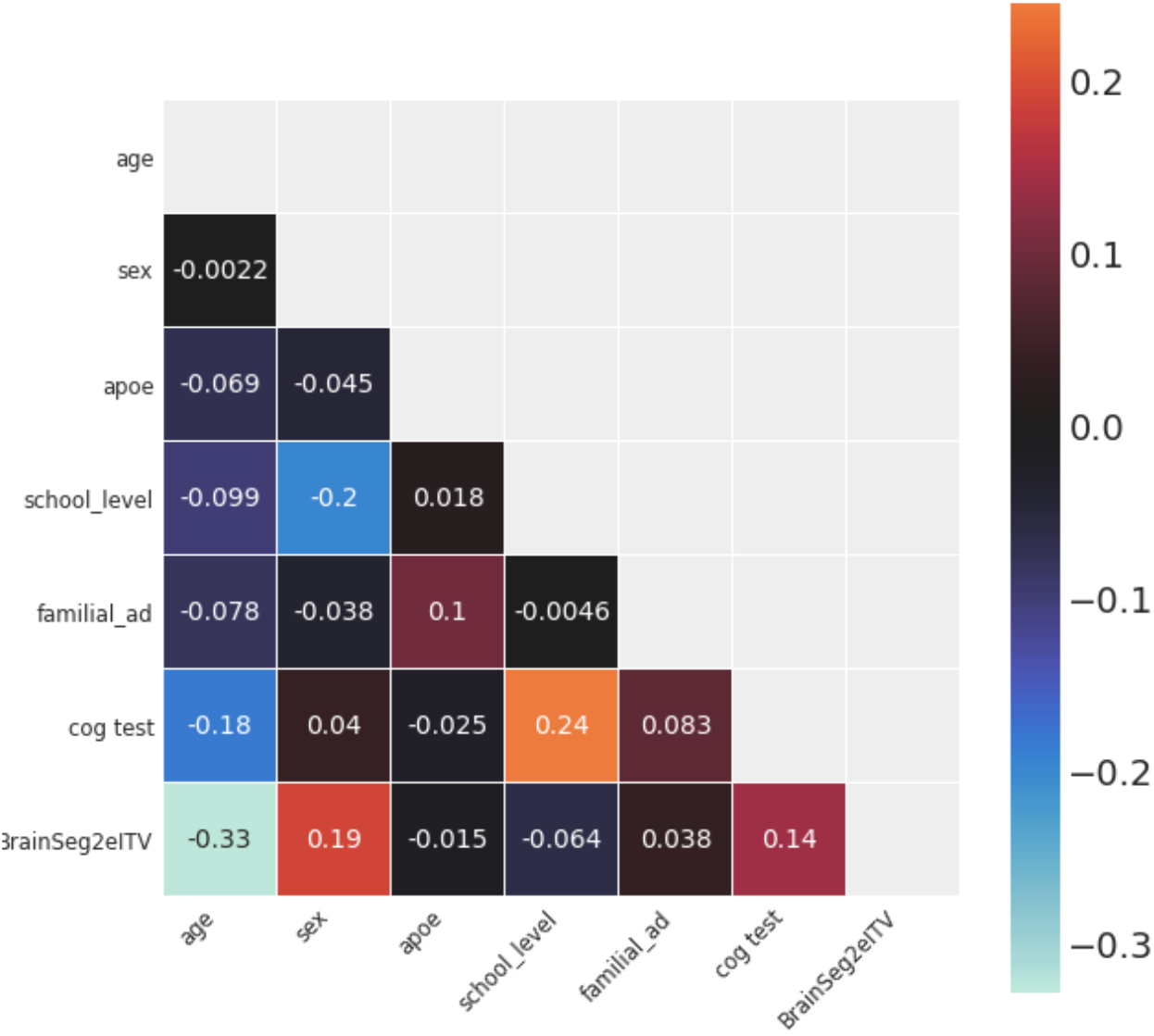
Correlation matrix of the variables used in the study. The variable of interest, brain volume to intracranial volume (Brain2ICV), is depicted in the last row. Age shows the strongest linear correlation with Brain2ICV (*ρ* = −.33) with Brain2ICV, followed by Sex (*ρ* = .19) and Memory test score (*ρ* = .14). School level, APOE and Family history of AD show no correlation with Brain2ICV.

### ii. Causal Analysis

Here we will find the answers to the two questions posed in Section II. To answer *Q1-Is there a direct causal relationship between Sex and Brain2ICV?-* we proceed by studying the difference in Brain2ICV between the male and female groups. We are thus, interested in the difference between the two groups rather than in the expected brain atrophy for each sex group which was already shown in Figure 3b. To compute this contrast, we use samples from the posterior distribution, that is to say we fit the model shown in Equation 2 to the data to have access to the posterior distribution of the difference or contrast between the male and female groups.

Table 3 shows the posterior distribution of the three parameters declared in Equation 2 (*μ*_1_, *μ*_2_, *σ*) and the posterior of the difference between the mean of brain atrophy between the male group and the female group or *μ*_1_ − *μ*_2_. The interpretation of the parameters in Table 3 is straightforward, the mean and standard deviation of the posterior distribution of Brain2ICV males is 0.697 ± 0.002 while for females is 0.708 ± 0.001. More importantly, the difference between the posterior distribution of the means shows that females get at old age with around 1% less atrophy than males, −0.011 ± 0.002. The high posterior density interval (HDI) is always negative, that is to say, when comparing the distribution of males and females, the area of the distribution Brain2ICV of females is larger than that of males. This confirms that sex plays a role in brain volume atrophy, with women getting into elderly age with slightly less brain loss volume relative to the maximum volume reached at a young age than men.

**Table 3:** The table shows the posterior distribution of the model’s parameters in Equation 2. The first two rows are the expected Brain2ICV (ratio between the brain volume (BV) at an elderly age and the intracranial volume (ICV) which works as a proxy of the maximum brain volume at a young age) in each sex group (1 for Male, 2 for Female), the third row is the standard deviation and the last row denotes the expected difference in Brain2ICV between (1) males and (2) females. The contrast, *μ*_1_ − *μ*_2_, is always negative which is indicative of less atrophy in women’s brains compared to men’s. This finding confirms previous studies in sex difference in the brain during aging, but here for the first time, via Bayesian statistical inference and posterior analysis rather than point estimates [Gamerman and Lopes, 2006], [Patil et al., 2010]. The High Posterior Density interval (HDI) is the shortest interval containing a give portion e.g 97% of the probability density. Note that HPI is not the same as a confidence interval, HDI is the probability of a variable having some value while a frequentist confidence interval contains or does not contain the true value of a parameter [Martin, 2018].

|  | mean | sd | HDI 3% | HDI 97% |
| --- | --- | --- | --- | --- |
| $\mu_1$ | 0.697 | 0.002 | 0.694 | 0.70 |
| $\mu_2$ | 0.708 | 0.001 | 0.706 | 0.71 |
| $\sigma$ | 0.027 | 0.001 | 0.026 | 0.028 |
| $\mu_1 - \mu_2$ | -0.011 | 0.002 | -0.014 | -0.007 |

Figure 5 shows graphically the posterior distribution and the sampling for the three unobserved variables in the model (*μ*_1_, *μ*_2_, *σ*). The variable Brain2ICV, *B_i_*, is the observed data and therefore we do not need to sample its values. On the left-hand of Figure 5, we show the kernel density estimation of the marginal distributions of each parameter and on the right hand we show the individual sampled values. The right-hand of Figure 5 shows

**Figure 5:**
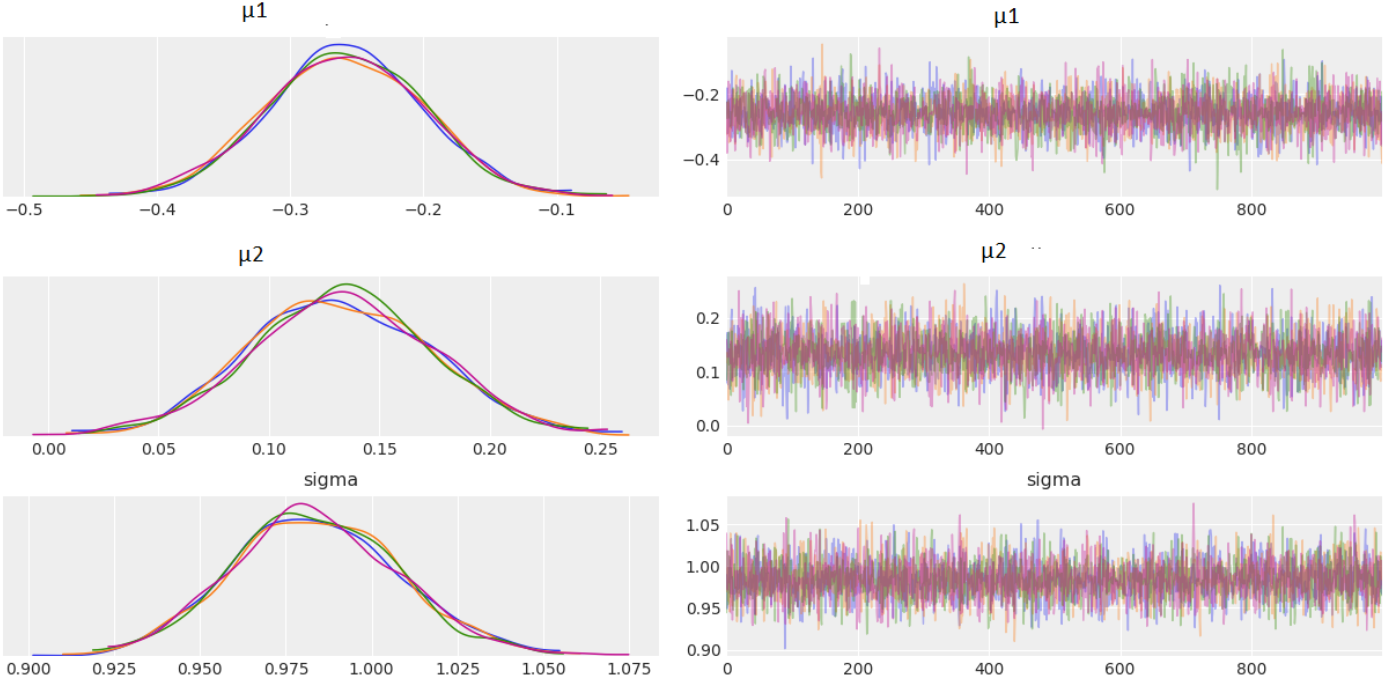
Posterior distribution (left-hand) and sampling (right-hand) for parameters *μ*_1_, *μ*_2_, *σ* (from top to bottom, Brain2ICV mean for men, Brain2ICV mean for women and Brain2ICV standard deviation) using PyMC3 [Salvatier et al., 2016]. On the left, kernel density estimation (KDE) plot for each of the 4 parallel chains plotted in different color run for each parameter *μ*_1_, *μ*_2_, *σ* (X-axis represents the value of the parameter and the Y-axis the Frequency). On the right, the individual sampled values at each step during the sampling for the four chains for each parameter(X-axis represents the sample number and the Y-axis the sample value). Males get into older age with more brain volume loss or less Brain to ICV ratio than women. The values have been normalized and standardized with mean 0 and standard deviation 1.

Males get into older age with more brain volume loss or less Brain to ICV ratio than women. The values have been normalized and standardized with mean 0 and standard deviation 1.

Figure 6a makes the comparison between the posteriors for male and female Brain2CV easy to visualize. As we can see, *μ*_1_ is negative and *μ*_2_ is positive with the HDI for the Male group slightly wider than the Female group. Since we have computed the posterior, we can use it to simulate data to assess the predictive quality of the model by comparing how consistently the simulated data match the observed data. Figure 6b shows the *posterior predictive checks* which allows us to evaluate the model by comparing the observed data and the model predictions (100 posterior predictive samples). The figure shows a good match between the mean and the variance of the simulated data and the actual data.

**Figure 6:**
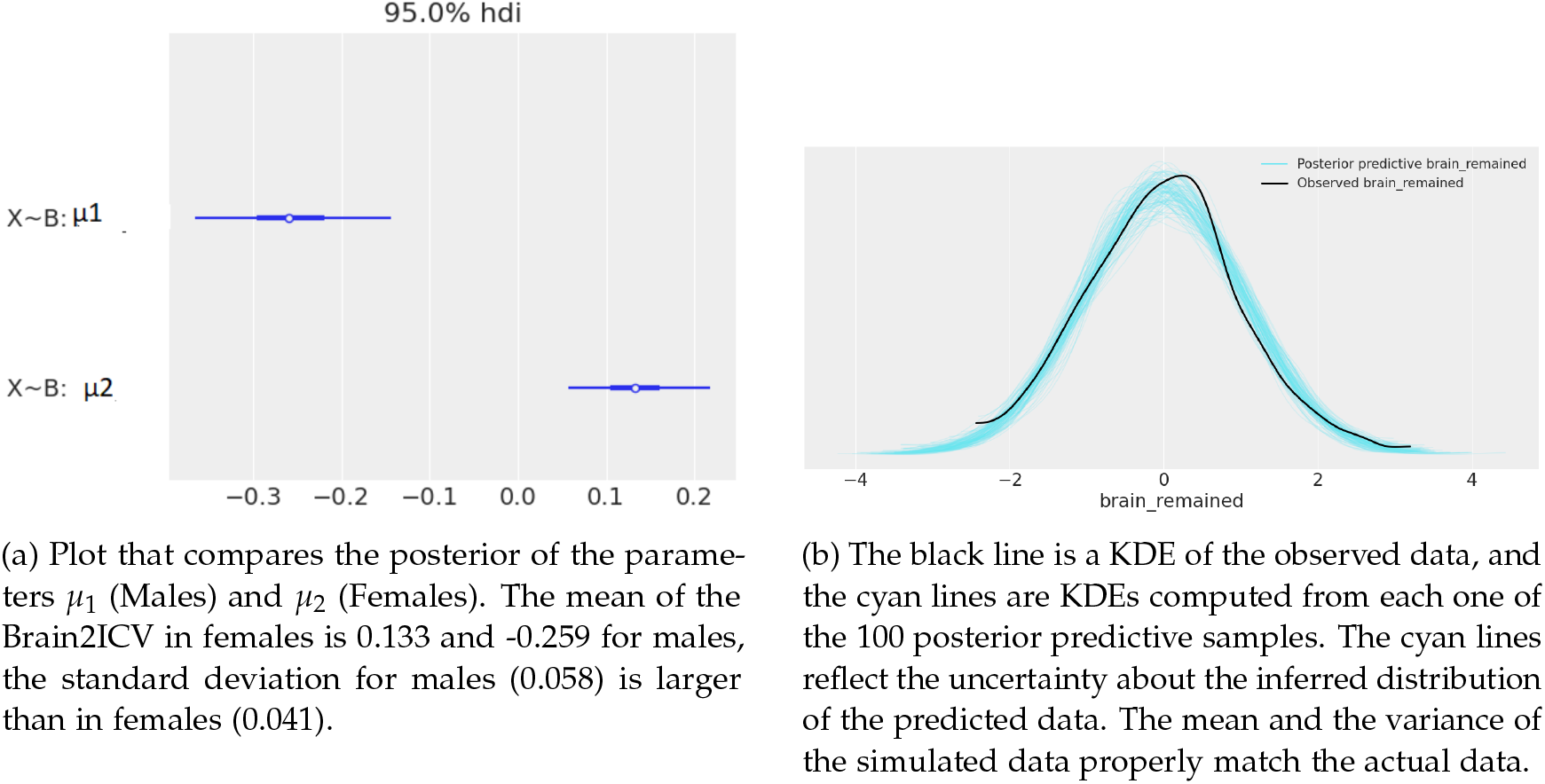
The black line is a KDE of the observed data, and cyan lines are KDEs computed from each one of the 100 posterior predictive samples. The cyan lines reflect the uncertainty about the inferred distribution of the predicted data. The mean and the variance of the simulated data properly match the actual data.

Now we are in a position to answer affirmatively Question 1:

> **Q1: Is there a direct causal relationship between brain atrophy -calculated as the ratio between brain size at an elderly age and maximum brain size at a young age- and sex? Yes. According to the Causal Diagram shown in Figure 2 Age has a direct effect on Brain2ICV. The expected difference or contrast between a female and a male in the sample shows that the expected Brain2ICV is larger for females.**

We address next *Q2* or *Is there additional value for Brain2ICV in knowing Sex if we already know Age?* The system of equations described in Equation 3 defines a multiple regression aimed at inferring the influence of Age on Brain2ICV while controlling for Sex and is represented in Equation 4 for convenience.

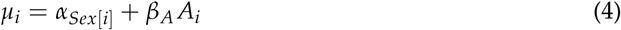

Table 4 shows the posterior distribution (mean standard deviation and HDI) of the multiple regression parameters declared in Equation 3 (*γ*_1_, *γ*_2_, *β*_*A*_, *σ* not shown for easiness). The first two rows denote the linear relationship between Brain2ICV and each sex group (1 for Male, 2 for Female), and the last row the linear relationship between Brain2ICV and Age. Each parameter in Table 4 controls the slope of the linear relationship and thus can be interpreted as the change in the outcome (Brain2ICV) per unit change in the predictor variables (Age and Sex). Thus, one standard deviation change in Age produces a change of one-third decrease in the standard deviation in Brain2ICV.

**Table 4:** Posterior distributions of the model’s parameters in Equation 3. Posterior of the average Brain2ICV for males *μ*_1_, posterior of the average Brain2ICV for females *μ*_2_ and Posterior of the age coefficient *β*_*A*_. The *β*_*A*_ coefficient describes a negative influence of the Age predictor on the outcome, Brain2ICV (negative mean). Similarly, there is a negative association between being Male and Brain2ICV. Thus, both Age and Masculine Sex are negatively associated with brain preservation at old age, computed as the ratio between the estimated brain volume (BV) and the estimated total intracranial volume (ICV). The Female Sex group, on the other hand, is positively associated with brain preservation.

|  | mean | sd | HDI 3% | HDI 97% |
| --- | --- | --- | --- | --- |
| $\mu_1$ | -0.266 | 0.052 | -0.369 | -0.177 |
| $\mu_2$ | 0.138 | 0.037 | 0.070 | 0.208 |
| $\beta_A$ | -0.33 | 0.031 | -0.392 | -0.277 |

Based on the results shown in Table 4 it is possible to graphically qualify, both in direction and magnitude, the weight that Sex and Age have separately on Brain2ICV. Figure 7 represents the mean and the standard deviation of the posteriors in the multiple regression shown in Table 4. The expected change per unit in Brain2ICV for elderly males is −0.266 and 0.138 for females. The expected change per unit in Brain2ICV for Age is −0.33, that is, an increase of one standard deviation in age reduces brings down the Brain2ICV by one third.

**Figure 7:**
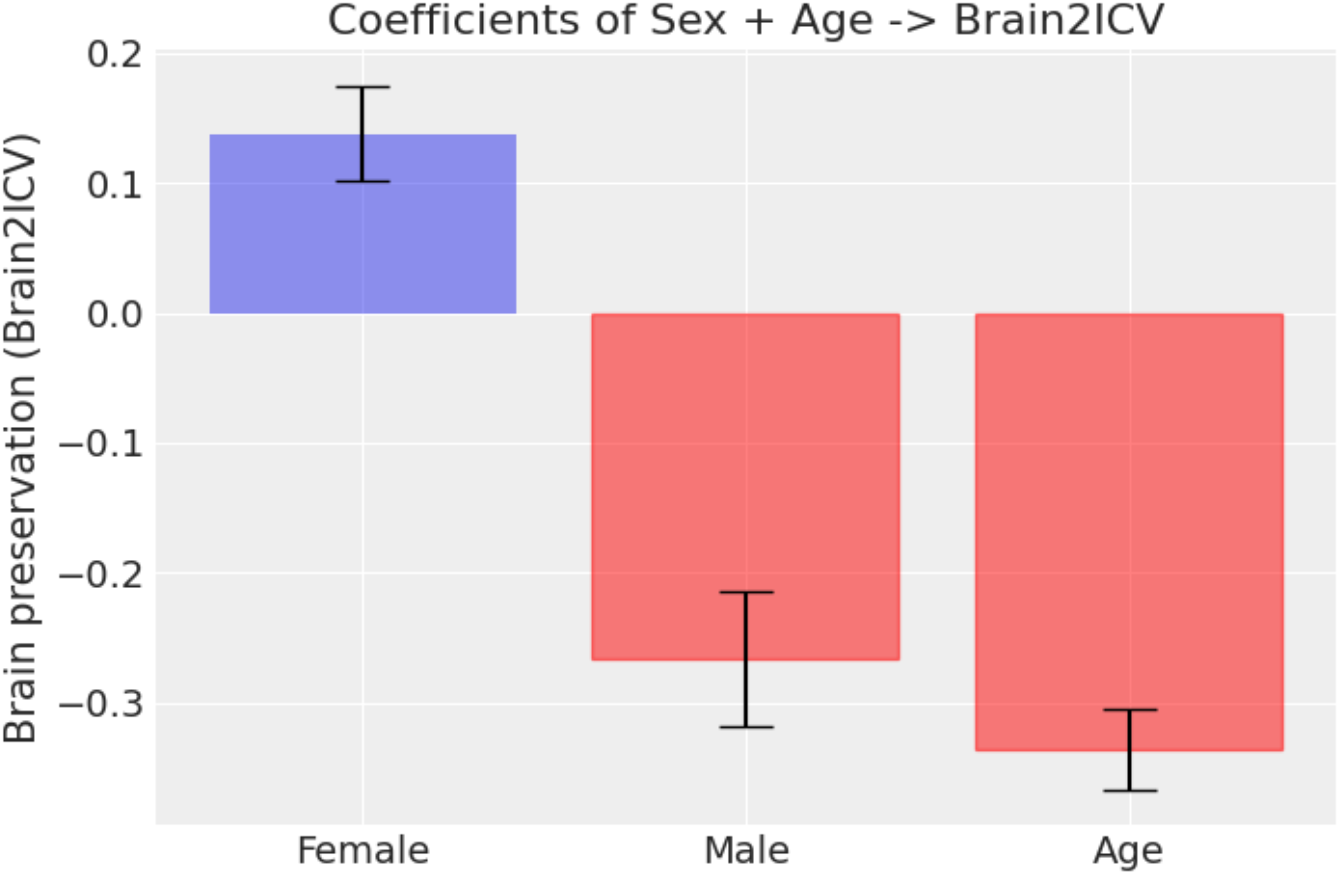
Direction and magnitude(*μ* and *σ*) of the posterior distribution of the parameters in the multiple regression model (Equation 4). Age and Male Sex are both negatively associated with brain preservation in older age, being Age, the stronger association. Female Sex is, on the other hand, positively associated although with a smaller effect than both Male Sex and Age.

Finally, Figure 8 allows to visually inspect the results of the multiple regression of the effect of both Age and Sex on Brain2ICV, *Age* + *Sex* → *Brain2ICV*, compared with the simple linear regressions *Sex* → *Brain2ICV*, and *Age* → *Brain2ICV* (shown in the Supplementary Materials). The main piece of information to extract from the figure is that once we know one of the two predictors, knowing the other one does not provide additional information about Brain2ICV. For instance, *β*_*A*_ has a very similar mean and standard deviation in the bivariate case (green bar) as in the multivariate regression case (blue bar). Or more easily put, if we know the Age, there is no additional gain in knowing also the Sex if we care about inferring Brain2ICV. Accordingly, we can answer Q2 in the negative.

> **Q2: Is there additional value for Brain2ICV in knowing Sex if we already know Age? No. Once we know the age of a subject knowing also her age conveys little information in predicting her Brain2ICV.**

**Figure 8:**
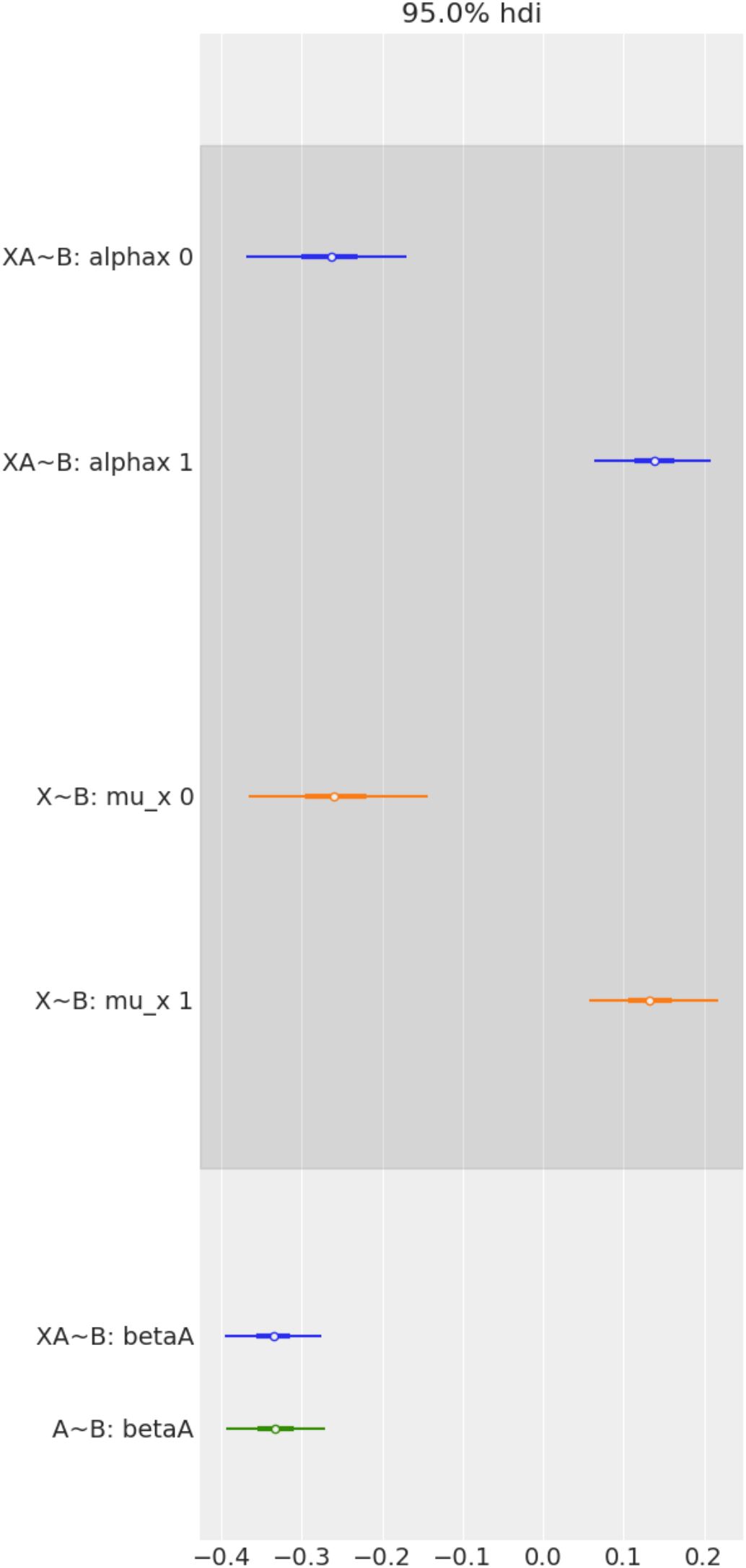
Forest plot to compare the effect of Age and Sex on Brain2ICV, separately and together in multiple regression. From top to bottom, the first two bars (in blue) represent the contrast in the multiple regression model, *Age + Sex* → *Brain2ICV*. The two middle bars (in orange) represent the posterior of Sex in the simple linear regression model *Sex* → *Brain2ICV*. The blue bottom bar depicts the posterior of Age in the multiple regression model, *Age + Sex* → *Brain2ICV*, and the green bar the posterior of Age but this time for the simple regression model, *Age* → *Brain2ICV*. Once we know Age (green bar at the bottom) there is no additional information in knowing also Sex (blue bar at the bottom) because the mean and the uncertainty remain mostly unchanged. Likewise, if we know Sex (orange bars) there is no significant information gain in also knowing Sex (blue bars at the top).

## IV. Discussion

The goal of this work is to study how brain atrophy is affected by factors such as age, APOE gene, sex, or school level among others. The study of brain volume loss at old age relative to young age requires, at least in principle, to have available MRI scans performed at either young and old age. There is, however, a way to address this problem having, as it is our case, only MRI scans at old age (70 years old or older). We compute the total brain loss of elderly subjects as the ratio between the estimated brain volume (BV) and the estimated total intracranial volume (ICV), called Brain2ICV for short. Thus, having only one MRI it is possible to ascertain the diminution of brain volume within the cranium relative to brain volume at a young age. We conceptualize the brain as a dynamic system inside a fossil container which sets the upper limit of brain volume. Accordingly, Brain2ICV represents the percentage of the volume occupied by the brain (BV) inside the cranium (ICV) which is used as a proxy of the maximum brain volume reached at a young age.

We find out that for the variables considered in this study -Age, APOE, Family history of AD, Sex, School Level- only Age and Sex affect Brain2ICV. Age is as expected, negatively associated with Brain2ICV, the older the brain gets the smaller is the ratio between the brain volume and the intracranial volume. More interestingly, we find that Sex plays a role in brain atrophy with Females having on average 1% larger Brain2ICV than males. In order of importance, age is the most important factor for brain volume loss, followed by masculine sex, and lastly feminine sex which is protective (Figure 7).

Methodologically speaking, this work departs from the approach of comparing differences between groups via point estimates and statistical testing. The pitfalls and difficulties associated with the statistical testing approach have been described both abundantly and convincingly [Association et al., 2016], [Benjamin et al., 2018], [Wasserstein et al., 2019] and we will therefore do not linger more on this point. Here, we follow a Bayesian approach to estimate posterior probability distributions rather than point estimates. It is worth reminding that under the Bayesian outlook, probabilities are tools to quantify uncertainty [Jaynes, 2003], [Gomez-Ramirez and Sanz, 2013]. Thus, we use probability distributions to summarize the entire plausibilities of each possible value of the parameter defined in the model. For example, the posterior distribution of the mean Brain2ICV in the female group entirely lies on the positive side, that is, we are certain that female sex and Brain2ICV are positively associated, while the opposite occurs for males, the posterior fully lies on the negative side which tells us that masculine sex and Brain2ICV are negatively associated.

The two questions addressed here: *Do age affects brain atrophy?* and *Which factors are causally connected with the loss of brain volume found in old age, sex, years of school, others?* are essentially causal inquiries and therefore cannot be responded to with correlations and data alone. We must postulate first a causal model which can be seen as the generator of the observed data. The causal diagram is postulated in Figure 2 and discussed in depth in Section ii.

The final goal of our methodological undertaking is no other than to achieve a causal understanding of the factors at play in the variability of brain volume loss in aging. To state that age causes aging is a platitude. However, behind this innocuous statement hides one of the defining scientific challenges of our time, Is aging inevitable? and Can we devise interventions aimed at modifying the aging process? Since we lack a theory of aging, what causes aging and how it progresses is ridden in uncertainty.

There are at least two major reasons for the lack of attention that causal reasoning has received in the scientific literature. One is historical and obeys to reasons not other than the personal preferences of leading scientists. As magisterially recounted by Judea Pearl in [Pearl and Mackenzie, 2018], causality was deliberately removed from statistics by Karl Pearson who considered cause and effect as animistic and unscientific concepts to be replaced by Contingency Tables which in Pearson’s mind were “the ultimate statement of the scientific description between two things” [Pearson, 1892]. The second and most important reason for the neglect of causality is that correlations, contrary to causal conclusions, do not require a controlled experiment and are therefore easy to compute.

The algebra created by Pearl, *do-calculus* [Pearl, 2009], [Pearl et al., 2016], can be treated as an extension of probability theory to investigate causal statements where before was only possible to perform observational analysis. While correlations have proved to be an extraordinarily successful tool to quantify pairwise relationships, correlations are lacking in situations where variables cannot be isolated. In such a scenario, we need to understand how the different variables interact with each other, which entails incorporating the direction in the association between two variables. A variable may cause another and this cannot be accounted for with correlations which are by definition symmetric.

We hope that the utility and promise of causal analysis have been rendered clear at this point. It is however worth recalling that causal analysis requires to postulate upfront the process that generates the data, that is to say, it requires the modeler’s input. The identification of causal associations will be consequently dependent on the model’s complexity. For example, in our case since we are interested in the effect of Sex and Age on Brain2ICV and both predictors are conditional independent, the causal analysis is fairly simple. Formally and in the language of *do-calculus*, Brain2ICV is a collider, *X* → *Y* ← *Z* (Sex(X), Age(Y), Brain2ICV(Z)). In a collider situation, conditioning on, for example, Z could induce a statistical association between X and Y misleading us into thinking that Age changes with Sex which is not the case. This is addressed in the multiple regression model (System of Equations 3) where we quantify each effect -Age on Brain2ICV and Sex on Brain2ICV- to find that the variables are conditionally independent *Y* ⫫ *X*|*Z*, (Y is not associated with X, after conditioning on Z) and *Y* ⫫ *Z*|*X* (Y is not associated with Z, after conditioning on X). Importantly, the same methodological framework can be used in more convoluted analysis, for example, the effect of Brain2ICV, Age, Sex, and School Years on Memory Test Scores (Figure 2). No direct causal path from Age to Memory or from Brain2ICV to Memory exists, thus the associations are in reality spurious and caused by the influence of variable(s) not included in the model (Supplementary Material).

This study is not without limitations. First, we are using whole-brain segmentation data without differentiating between brain tissues or brain anatomical structures. Second, in positing as we do a causal model, we are aiming at the causal mechanisms involved in brain aging which a complex biological process that lacks a coommonly accepted definition [Viña et al., 2007]. Second, the causal diagram postulated in Figure 2 may be incomplete as it is the case, for example, in the causal relationships between variables related to memory. Nevertheless, our focus on the direct effect on brain volume atrophy makes both limitations not particularly concerning. Although a more granular data acquisition of brain anatomy would convey information that is missing in this study, the approach used here makes it uniquely possible to investigate the shrinkage of the brain using only one measurement (a single MRI). Since the brain is contained within the scaffolding of the cranium, we can estimate the total brain volume loss taking the intracranial volume as the asymptotic volume. This method is impractical for structures inside the cerebrum since we would be missing the fossil-like container that the cranium provides for the entire organ.

Part of the novelty and interest of the study relies upon its methodological underpinning, which departs from point estimates and linear associations between variables. We use Bayesian probabilistic programming to study in a principled way causal inference, combining the flexibility of Bayesian probability and the applicability of sampling theory in a coherent decision theoretical framework. Although it is beyond reach to verify the validity or completeness of causal diagrams, a causal diagram together with the appropriate algebra (do-calculus) enable causal reasoning, paving the way from hypothesis testing and correlations to interventions and counterfactual reasoning, reaching thus the proper epistemological scaffolding required to capture the structure that generates events in nature.

## V. Data availability

The data set can be downloaded downloaded from https://github.com/grjd/causalityagingbrain.

## VI. Code availability

The code and data set are available at: https://github.com/grjd/causalityagingbrain.

## VII. Acknowledgments

The authors would like to thank the participants in the present study. We acknowledge funding from *Ministerio de Ciencia, Innovación y Universidades* (CONNECT-AD) RTI2018-098762-B-C31 and Structural Funds ERDF (INTERREG V-A Grant: 0348CIE6E).

